# Interplay between receptor binding, immune escape and protein stability determines the natural selection of SARS-CoV-2 variants

**DOI:** 10.1101/2021.05.23.445348

**Authors:** Vaibhav Upadhyay, Alexandra Lucas, Sudipta Panja, Ryuki Miyauchi, Krishna M.G. Mallela

**Affiliations:** Department of Pharmaceutical Sciences, Skaggs School of Pharmacy and Pharmaceutical Sciences, University of Colorado Anschutz Medical Campus, Aurora, CO 80045

**Author notes:** Corresponding author Krishna Mallela, PhD, Department of Pharmaceutical Sciences, University of Colorado Anschutz Medical Campus, 12850 E. Montview Blvd, MS C238-V20, Aurora, CO 80045.

**Keywords:** protein stability, protein binding, protein function, protein structure, protein expression, COVID-19, SARS-CoV-2, ACE2, antibodies, variants

## Abstract

Emergence of new SARS-CoV-2 variants has raised concerns at the effectiveness of vaccines and antibody therapeutics developed against the unmutated wild-type virus. We examined the effect of 12 most commonly occurring mutations in the receptor binding domain on its expression, stability, activity, and antibody escape potential-some of the factors that may influence the natural selection of mutants. Recombinant proteins were expressed in human cells. Stability was measured using thermal denaturation melts. Activity and antibody escape potential were measured using isothermal titration calorimetry in terms of binding to ACE2 and to a neutralizing human antibody CC12.1, respectively. Our results show that variants differ in their expression levels with the two least stable variants showing lesser expression. Out of the 8 well-expressed mutants, only 2 (N501Y and K417T/E484K/N501Y) showed stronger affinity to ACE2, 4 (Y453F, S477N, T478I and S494P) have similar affinity, whereas the other 2 (K417N and E484K) have weaker affinity when compared to the wild-type. In terms of CC12.1 binding, when compared to the wild-type, 4 variants (K417N, Y453F, N501Y and K417T/E484K/N501Y) have weaker affinity, 2 (S477N and S494P) have similar affinity, and 2 (T478I and E484K) have stronger affinity. Taken together, these results indicate that multiple factors contribute towards the natural selection of variants, and all these factors need be considered to understand the evolution of the virus. In addition, since not all variants can escape a given neutralizing antibody, antibodies to treat new variants can be chosen based on the specific mutations in that variant.

## INTRODUCTION

Corona virus disease 2019 (COVID-19) pandemic has emerged as a global threat in December 2019. As of August 2021, it has infected over 218 million people and claimed 4.5 million lives (https://coronavirus.jhu.edu/map.html). The causative agent for COVID-19 is severe acute respiratory syndrome coronavirus 2 (SARS-CoV-2), a single stranded RNA virus, that belongs to sarbecovirus subgenus of betacoronaviruses (1). It shares close genomic similarity to SARS-CoV (79% identity) and Middle East respiratory syndrome coronavirus (MERS-CoV, 50% identity) that were responsible for SARS outbreak in 2002 and MERS outbreak in 2012 respectively (2-6). These viruses are thought to have originated in bats and transmitted to other mammals including humans (7).

Both SARS-CoV and SARS-CoV-2 enter the human host through interaction of its spike protein with angiotensin-converting enzyme 2 (ACE2) present on the membrane of host epithelial cells (8-10). Specifically, the receptor binding domain (RBD) of the spike protein binds with ACE2 and thus is a major determinant of the viral infectivity and evolution (11-13). The viral evolution through accumulation of mutations in SARS-CoV-2 is slower than known for other RNA viruses like HIV and influenza (14-16). Still, SARS-CoV-2 variants pose a major challenge for devising measures to counter the virus threat, as new variants continue to emerge, some of which are believed to be more infectious than the wild-type virus (17-19). Among all the mutations happening in the viral genome, mutations in RBD are considered to play a significant role in infectivity due to its role in ACE2 binding. Most of the neutralizing antibody response of the host is generated against RBD (20,21), because of which RBD is also a major target for most of the therapeutic antibodies developed (22-24). Mutations in RBD are predicted to dictate the emergence of escape mutants and shape the evolutionary path of the virus through the process of natural selection that would favor the mutants that could evade the antibody response. Apart from this, RBD alone or as part of spike protein is also used as antigen in many prospective vaccines (25-35). The emergence of mutations in RBD is considered to have an impact on the effectiveness of these vaccines. Lower efficacy of some of the vaccines was reported against the emerging Variants of Concern (VOC) that include Alpha, Beta and Gamma variants along with reports of new variants escaping the antibodies approved for emergency use (36-39).

In this work, we examined the effect of variants on RBD protein expression, stability, its binding to ACE2, and antibody escape using a naturally occurring human neutralizing antibody CC12.1. We hypothesize that all these factors act in conjunction and can determine the virus evolution and natural selection of new variants. To test, we selected 12 most frequently occurring mutations in RBD as of January 2021 (https://www.gisaid.org/hcov19-mutation-dashboard). These include 9 single site mutations K417N, N439K, Y453F, S477N, S477I, T478I, E484K, S494P and N501Y (Alpha variant), a double mutant (E484K/N501Y) and two triple mutants (K417N/E484K/N501Y (Beta variant) and K417T/E484K/N501Y (Gamma variant)). Location of these residues in RBD bound to ACE2 and CC12.1 are shown in Figs. 1A and 1B. All these mutating residues are in the receptor binding motif (RBM) of RBD which interacts with ACE2 receptor. Our results presented below indicate that multiple factors contribute towards the natural selection of variants, and all these factors must be considered to understand the natural selection of SARS-CoV-2 variants.

**Figure 1.**
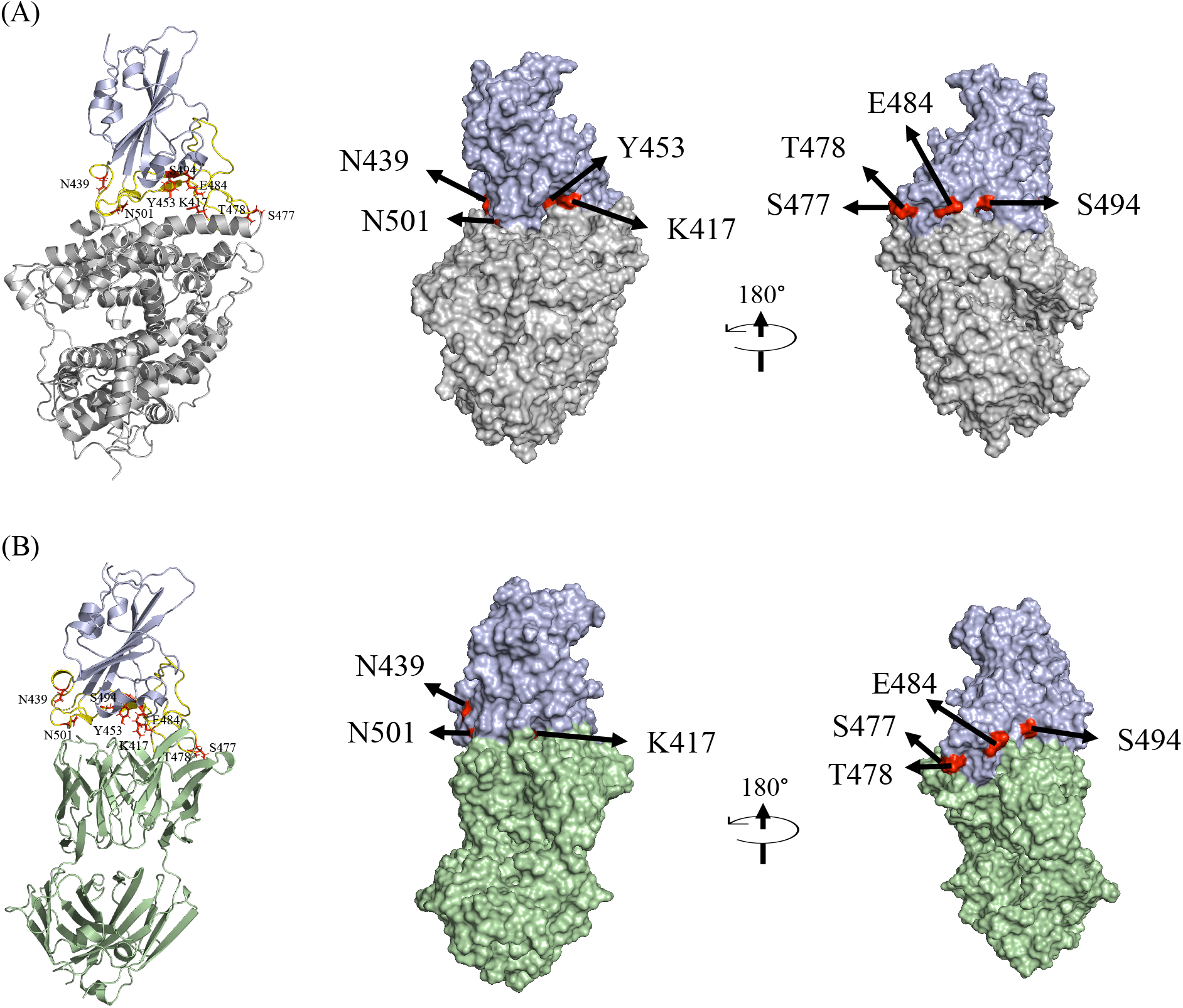
Structures of SARS-CoV-2 RBD (colored blue) interacting with (A) ACE2 (colored gray; PDB ID 6m0j) and (B) CC12.1 Fab (colored green; PDB ID 6xc2), showing the most frequently mutating residues in RBD-K417, N439, Y453, S477, T478, E484, S494 and N501 (colored red). RBM is shown in yellow color. The single mutants of RBD used in this study were K417N, N439K, Y453F, S477N, T478I, E484K, S494P and N501Y (Alpha variant). A double mutant (E484K/N501Y) and triple mutants corresponding to Beta variant (K417N/E484K/N501Y) and Gamma variant (K417T/E484K/N501Y) were also used. The position of Y453 is not visible in the surface view of RBD interacting with CC12.1 Fab as it is buried at the interface.

## RESULTS

### RBD mutations affect protein expression

Protein expression was performed in human embryonic kidney (HEK) cells, in which the protein transport through the secretory pathway, the post-translational modifications and quality control mechanisms ultimately decide the secreted protein levels (40). Expression in HEK cells closely match the natural infection scenario, where the virus uses the host cell machinery to synthesize its structural proteins. The protein expression levels have a direct bearing on the yield of the viruses, as the virus yield is directly proportional to the amount of the proteins available for virus assembly (41). The infectivity of a particular variant can thus be dependent on the protein expression levels. It should be noted however, that this study only represents the expression level of RBD and in natural scenario the expression level of the complete spike protein would ultimately decide the virus yield and thus infectivity. The relative expression of the wild-type RBD along with the mutants were compared using SDS-PAGE after 3 days of expression (Figs. 2A and 2B). Among 9 single site mutants, 2 mutants (N439K and S477I) did not express very well and appreciable amount of protein needed for binding studies could not be obtained for these mutants. The levels of expression were also low for 2 other single site mutants T478I and E484K. Expression levels comparable to or higher than the wild type was obtained for 5 single site mutants K417N, Y453F, S477N, S494P and N501Y. Apart from the single site mutations, clone carrying double mutations E484K/N501Y did not express. The clone carrying triple mutations (K417N/E484K/N501Y) corresponding to the Beta variant also could not be expressed, but the other clone carrying triple mutations (K417T/E484K/N501Y) corresponding to the Gamma variant showed high expression. These results suggest that RBD mutations can impact the overall protein expression levels. Similar mutation effects on the expression of the complete spike protein might exist, which can affect the virus yield and infectivity.

**Figure 2.**
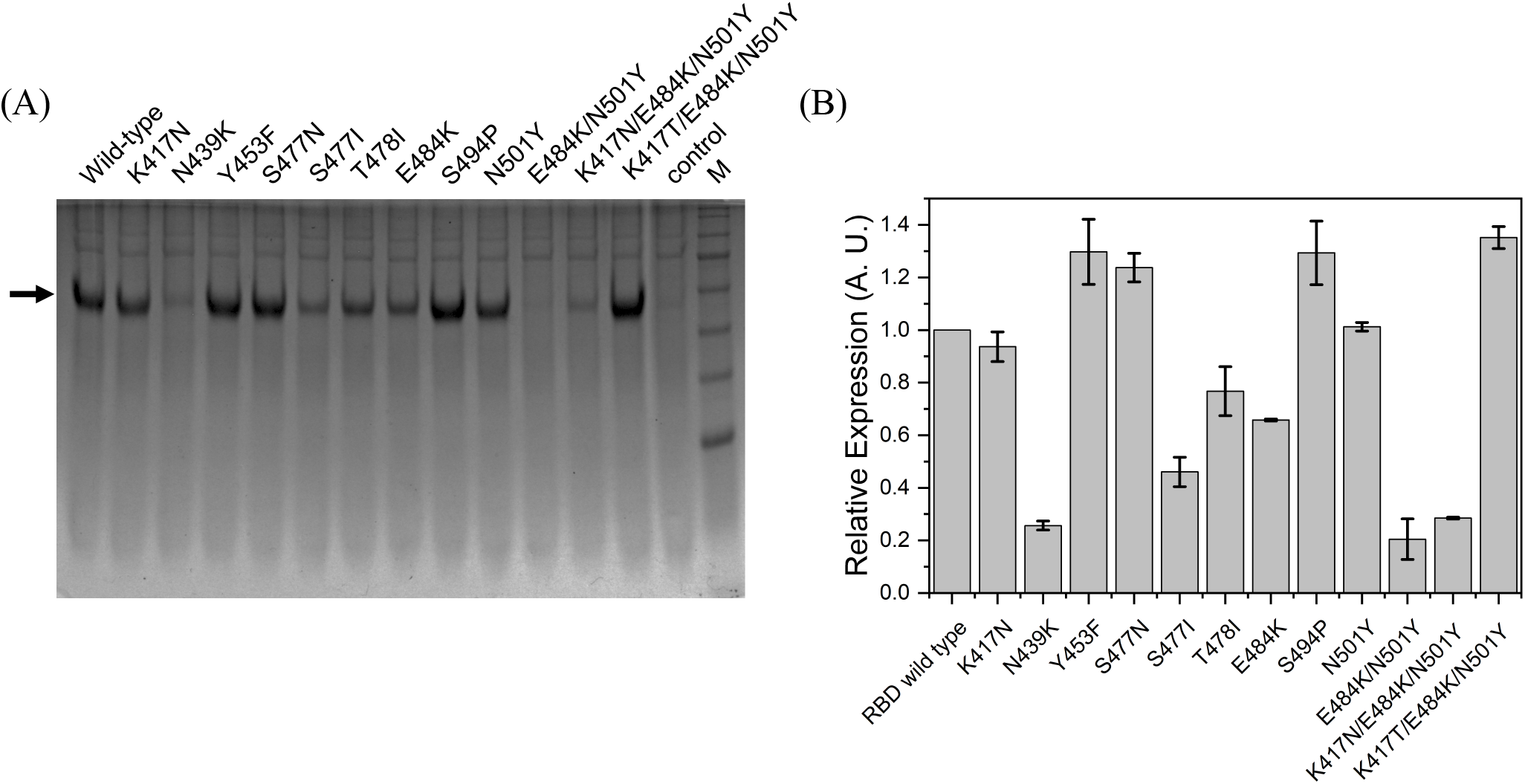
Comparison of relative expression of RBD and its mutants. (A) SDS-PAGE showing relative amounts of expressed RBD and its mutants. M represents molecular weight markers (From top to bottom: 180, 130, 100, 70, 55, 35 and 25 kDa) (B) Relative expression of mutants quantified from the band intensities in SDS-PAGE in panel A.

Because of protein expression constraints, further studies on the mutant proteins were carried out on 7 single site mutants (K417N, Y453F, S477N, T478I, E484K, S494P and N501Y) and the triple mutant K417T/E484K/N501Y, which were purified to homogeneity along with ACE2 and CC12.1 single chain variable fragment (ScFv) (Figs. 3A and 3B).

**Figure 3.**
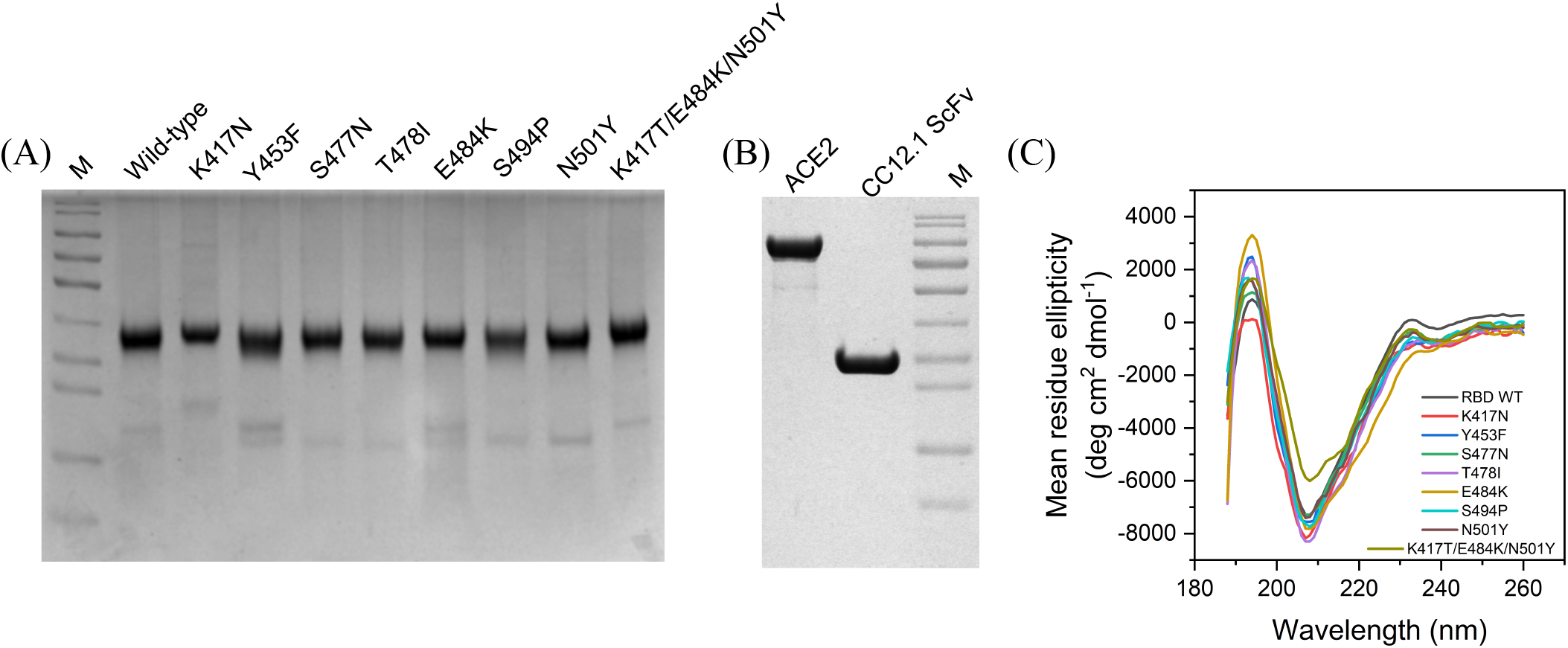
(A) Purified RBD and its mutants (B) ACE2 and CC12.1 ScFv. M represents molecular weight marker (From top to bottom: 180, 130, 100, 70, 55, 35, 25, 15 and 10 kDa). (C) Comparison of secondary structures of RBD and its mutants using far-UV CD spectroscopy. Table 1 lists the proportion of various secondary structures when the spectra were deconvoluted using BeStSel software.

### Mutations do not significantly affect the global protein structure

Most of the random mutations in proteins do not affect the protein structure and are thus considered neutral. Some mutations though, can bring significant structural changes, and can have either a beneficial or a deleterious effect on virus fitness. In the absence of other selection pressures, the deleterious mutations are lost but the beneficial mutations get selected and prevail. Here we investigated the effect of 8 frequently occurring mutations on the RBD structure using far-UV circular dichroism (CD) spectroscopy (Fig. 3C). The CD spectra for the wild-type protein showed a major negative band at 208 nm and a positive band at 192 nm. The CD spectra was similar to previously reported spectra for wild-type RBD (42). The far-UV CD spectra of all 8 mutants were similar to that of the wild-type protein with most significant difference observed for the Gamma variant. Deconvolution using BeStSel web software (43) reveal a low percentage alpha-helix along with high beta-sheet content for all the variants. The proportion of the random coil structure is substantial for RBD accounting for about 50% (Table 1), primarily originating from the RBM (Fig. 1). These CD results suggest that the mutations do not significantly affect the global structure of RBD, and all mutants adopt a similar fold like the wild-type RBD. This is also consistent with the soluble expression of variants (Fig. 3A). Proteins that are expressed as secretory proteins in general adopt well-folded structures. This shows that only those mutants that do not have any deleterious effect on the protein structure are getting naturally selected. Since software used for deconvoluting CD spectra in general accounts only for regular secondary structures, minor differences in the CD spectra of the triple mutant K417T/E484K/N501Y could be due to changes in short, irregular and non-repeating secondary structures most probably in the RBM region.

**Table 1.** Proportion of the secondary structural elements calculated by deconvolution of far-UV CD spectra of RBD and its mutants using BeStSel software (43).

| RBD variant | Secondary structure |  |  |  |  |
| --- | --- | --- | --- | --- | --- |
|  | Helix | Antiparallel | Parallel | Turn | Others |
| Wild-type | 13.7 | 25.0 | 0.0 | 13.9 | 47.3 |
| K417N | 12.8 | 17.9 | 4.4 | 14.1 | 50.7 |
| Y453F | 15.3 | 19.2 | 1.5 | 13.9 | 50.2 |
| S477N | 13.5 | 22.0 | 2.0 | 13.2 | 49.3 |
| T478I | 13.8 | 17.8 | 3.2 | 13.3 | 51.9 |
| E484K | 15.7 | 16.3 | 4.0 | 13.4 | 50.6 |
| S494P | 14.1 | 21.4 | 2.1 | 13.3 | 49.1 |
| N501Y | 11.4 | 22.9 | 2.8 | 13.0 | 49.9 |
| K417T/E484K/N501Y | 10.4 | 24.3 | 2.9 | 13.1 | 49.3 |

### Most RBD mutants have similar thermal stability as that of wild-type protein

Protein stability can be an important factor in protein evolution (44). More stable proteins can accommodate wide range of mutations and determine the evolvability of the proteins (45). It is thus important to determine the stability of protein to be able to determine its evolutionary path. The stability of the wild-type RBD and its mutants were assessed with thermal denaturation melts using far-UV CD spectroscopy (Fig. 4). Thermal denaturation of the wild-type protein shows a co-operative transition with slopy native and denatured baselines. The unfolding transition could be fitted well to a two-state model (Eq. 1 in Experimental Procedures), giving T_m_ (midpoint temperature of thermal denaturation) value of 56.1 ± 0.7 °C (Table 2). The thermal denaturation curves of the mutant proteins were similarly obtained and were found to be cooperative, which fitted well to a two-state unfolding model. All the mutants showed similar T_m_ values as that of the wild-type protein, indicating that none of the mutations are causing drastic changes in the RBD stability. Compared to other mutants, T478I and E484K showed slightly lesser thermal stability. The same two mutants also showed lesser expression levels (Fig. 2B).

**Figure 4.**
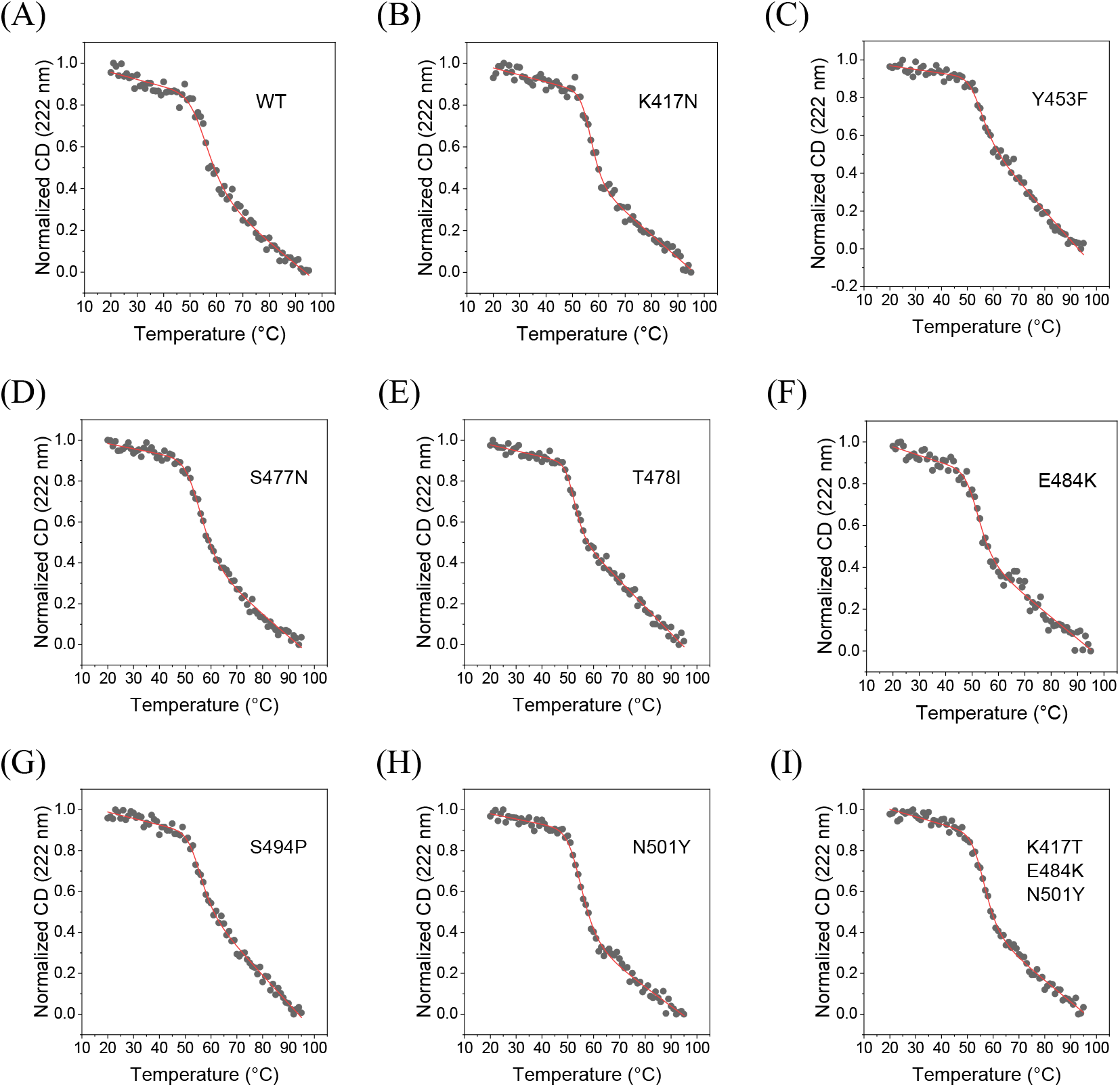
Thermal denaturation melts of RBD and its mutants obtained using far-UV CD spectroscopy. Panels A-I show the data for the wild-type RBD, single amino acid mutations K417N, Y453F, S477N, T478I, E484K, S494P, N501Y, and for the triple mutant K417T/E483K/N501Y, respectively. The solid lines show the fits to a 2-state unfolding equation (Eq. 1 in Methods section). Table 2 lists the T_m_ (midpoint meting temperature) and the ΔH (enthalpy change at Tm) values of RBD variants.

**Table 2.** T_m_ and **Δ**H values of RBD variants obtained from thermal denaturation curves using far-UV CD. Errors on ΔT_m_ were calculated using error propagation formulae (72).

| RBD variant | $T_m$ (°C) | $\Delta T_m$ (°C) | $\Delta H$ (kcal/mol) |
| --- | --- | --- | --- |
| Wild-type | $56.1 \pm 0.7$ | - | $-1.7 \pm 0.3$ |
| K417N | $56.9 \pm 0.3$ | $0.8 \pm 0.8$ | $-3.0 \pm 0.4$ |
| Y453F | $54.8 \pm 0.4$ | $-1.3 \pm 0.8$ | $-2.2 \pm 0.3$ |
| S477N | $55.8 \pm 0.3$ | $-0.3 \pm 0.8$ | $-1.9 \pm 0.1$ |
| T478I | $52.3 \pm 0.2$ | $-3.8 \pm 0.7$ | $-2.9 \pm 0.3$ |
| E484K | $52.3 \pm 0.5$ | $-3.8 \pm 0.9$ | $-2.0 \pm 0.3$ |
| S494P | $55.3 \pm 0.4$ | $-0.8 \pm 0.8$ | $-2.3 \pm 0.3$ |
| N501Y | $55.1 \pm 0.3$ | $-1 \pm 0.8$ | $-2.1 \pm 0.1$ |
| K417T/E484K/N501Y | $56.5 \pm 0.2$ | $0.4 \pm 0.7$ | $-2.4 \pm 0.2$ |

### Some but not all RBD mutants show increased binding affinity to ACE2

SARS-CoV-2 RBD mediates the interaction of the virus spike protein with the ACE2 receptor on the host cell surface and is thus, a major determinant of the viral entry into the host cell. Mutations in the RBD can impact its interaction with ACE2 and can have an important role in determining the infectivity of the virus, with higher affinity interactions contributing to increased infectivity. The binding interactions of the wild-type and mutant RBD proteins with ACE2 were investigated using isothermal titration calorimetry (ITC) (Fig. 5 and Table 3). ITC is particularly advantageous in terms of measuring protein-protein interactions in solution without any covalent modification of proteins. The wild-type RBD interacts with ACE2 in a stoichiometric ratio of 1:1, with a K_d_ value of 10.0 ± 3.1 nM and enthalpy of interaction (ΔH) of 11.8 ± 0.2 kcal/mol. The measured K_d_ value is consistent with previously published studies on ACE2-RBD interaction using surface plasmon resonance (SPR) with immobilized protein (46). All the mutants studied interacted with ACE2 in the same stoichiometric ratio of 1:1. Three of the 8 mutants, Y453F, T478I and S494P, did not show significant difference in their binding interaction with ACE2, with K_d_ and ΔH values similar to the wild-type protein (Table 3). For S477N mutant, K_d_ value was similar to that of the wild-type, but an increased ΔH value of 16.4 ± 0.2 kcal/mol was obtained, which may indicate increased interactions between RBD and ACE2 upon mutation. For two other mutants K417N and E484K, the K_d_ value obtained was higher than the wild-type protein (Table 3), indicating weaker affinity for ACE2. The corresponding ΔH values did not show any significant difference for K417N mutant but showed a decreased value of 9.5 ± 0.2 kcal/mol for E484K mutant. The other single site mutant N501Y corresponding to the Alpha variant and the triple mutant K417T/E484K/N501Y corresponding to the Gamma variant showed increased affinity for ACE2 binding with K_d_ values of 3.0 ± 2.1 nM and 1.6 ± 1.5 nM respectively when compared to the wild-type (Table 3). There was no difference in ΔH values for N501Y mutant and the triple mutant K417T/E484K/N501Y. These ITC results indicate that the ACE2 binding is an important but not the sole parameter determining the natural selection of the variants.

**Figure 5.**
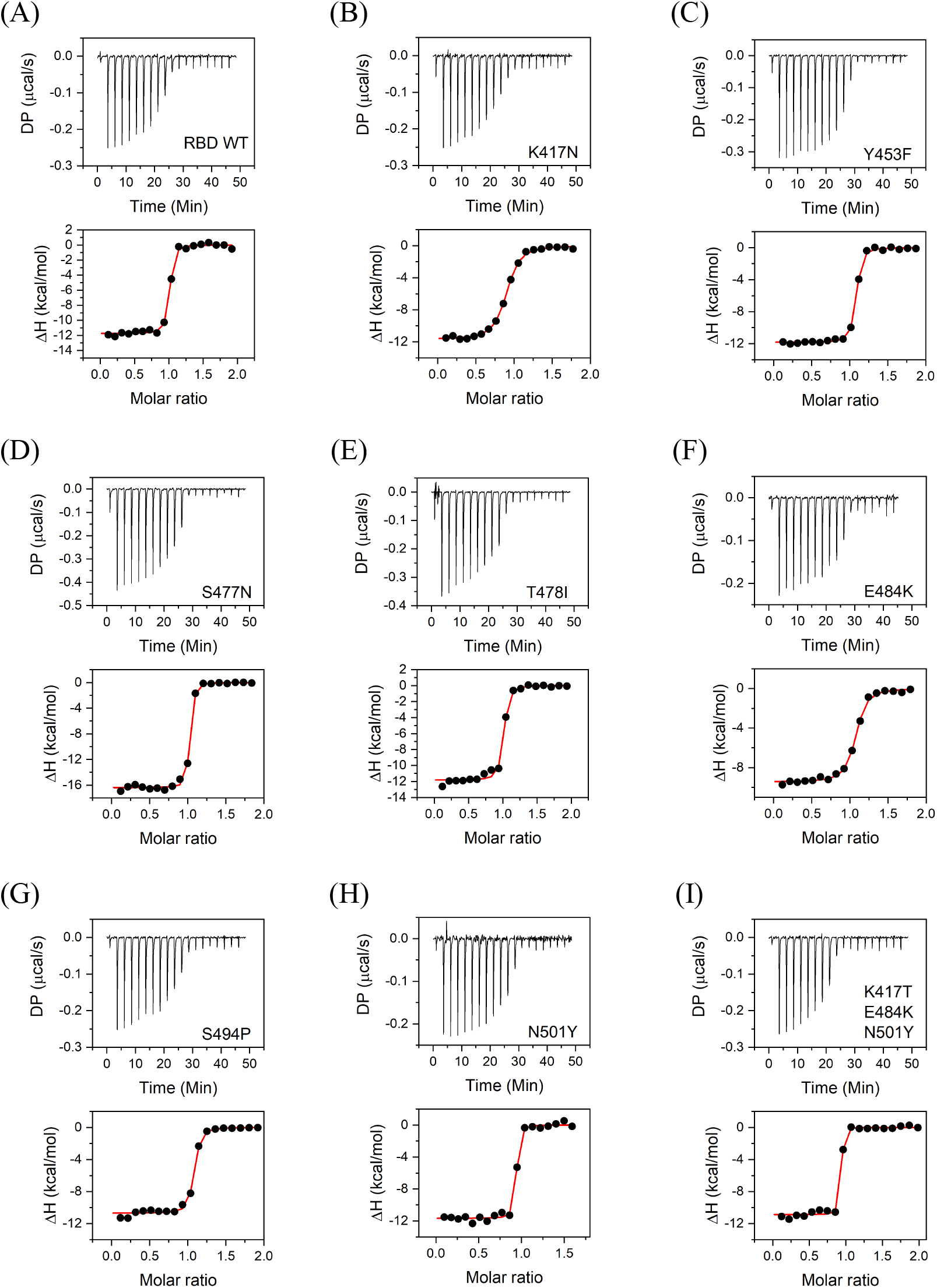
Binding of RBD and its variants to ACE2 studied using ITC. Panels A-I show the data for the wild-type RBD, single amino acid mutations K417N, Y453F, S477N, T478I, E484K, S494P, N501Y, and for the triple mutant K417T/E483K/N501Y, respectively. Top panels show the raw thermograms, and the bottom panels show the fit to the integrated heat curve. Table 3 lists the interaction parameters from the data fit.

**Table 3.** Interaction parameters obtained from binding of RBD variants to ACE2 probed by ITC. Errors on ΔG and -TΔS were calculated using error propagation formulae (72).

| **RBD variant** | **K<sub>d</sub>**<br>**(nM)** | **N** | **$\Delta H$**<br>**(kcal/mol)** | **$\Delta G$**<br>**(kcal/mol)** | **$-T\Delta S$**<br>**(kcal/mol)** |
| --- | --- | --- | --- | --- | --- |
| Wild-type | 10.0 ± 3.1 | 1.0 ± 0.0 | -11.8 ± 0.2 | -10.7 ± 0.2 | 1.0 ± 0.3 |
| K417N | 94.1 ± 9.5 | 0.9 ± 0.0 | -11.7 ± 0.1 | -9.4 ± 0.1 | 2.3 ± 0.1 |
| Y453F | 11.4 ± 1.7 | 1.0 ± 0.0 | -11.8 ± 0.1 | -10.7 ± 0.1 | 1.2 ± 0.1 |
| S477N | 5.0 ± 1.4 | 1.0 ± 0.0 | -16.4 ± 0.2 | -11.1 ± 0.2 | 5.3 ± 0.3 |
| T478I | 16.7 ± 5.6 | 1.0 ± 0.0 | -11.8 ± 0.2 | -10.4 ± 0.2 | 1.4 ± 0.3 |
| E484K | 51.5 ± 8.2 | 1.0 ± 0.0 | -9.5 ± 0.2 | -9.8 ± 0.1 | -0.3 ± 0.2 |
| S494P | 13.8 ± 3.5 | 1.0 ± 0.0 | -10.7 ± 0.2 | -10.5 ± 0.1 | 0.2 ± 0.2 |
| N501Y | 3.0 ± 2.1 | 0.9 ± 0.0 | -11.7 ± 0.2 | -11.4 ± 0.4 | 0.2 ± 0.5 |
| K417T/E484K/N501Y | 1.6 ± 1.5 | 0.9 ± 0.0 | -10.9 ± 0.2 | -11.8 ± 0.5 | -0.9 ± 0.6 |

### Some but not all RBD mutants show CC12.1 antibody escape

CC12.1 antibody is a monoclonal antibody isolated from the convalescent plasma of a COVID-19 survivor (47,48). This antibody represents the class of antibodies most elicited by SARS-CoV-2 infection and also in response to current vaccines against the wild-type. The circulating antibodies can play a major role in shaping the evolutionary path of RBD. The mutants that can escape the antibody recognition are naturally selected and represented more in the viral pool. We investigated the binding of the wild type RBD and its mutants to the ScFv of CC12.1 antibody through ITC (Fig. 6 and Table 4). The CC12.1 ScFv interacts with wild type RBD at a stoichiometric ratio of 1:1 with a K_d_ value of 28.0 ± 8.8 nM and ΔH value of 4.8 ± 0.1 kcal/mol. Compared to ACE2 interaction which is primarily enthalpically driven (Table 3), RBD interaction with CC12.1 has a significant entropic component (Table 4). Out of the 8 mutants studied, 2 single site mutations (S477N and S494P) did not impact the binding affinity of RBD towards CC12.1 ScFv. E484K and T478I mutants however showed increased affinity towards CC12.1 ScFv with a K_d_ value of 1.7 ± 4.6 and 5.8 ± 3.4 nM respectively (Table 4), which suggests that CC12.1 may be able to neutralize E484K and T478I mutants. Other 4 variants, single site mutants K417N, Y453F, N501Y (Alpha variant) and the triple mutant K417T/E484K/N501Y (Gamma variant), showed decreased affinity towards CC12.1 binding, with K_d_ values of 119 ± 50 nM, 827 ± 146 nM, 63 ± 22 and 433 ± 95 nM respectively, representing escape from CC12.1. Another interesting observation is the decrease in ΔH value for K417N and Y453F mutants, indicating decreased strength of interactions between the mutants and CC12.1. These results show that antibody escape can be a very important parameter that shapes virus evolution and natural selection of mutants.

**Figure 6.**
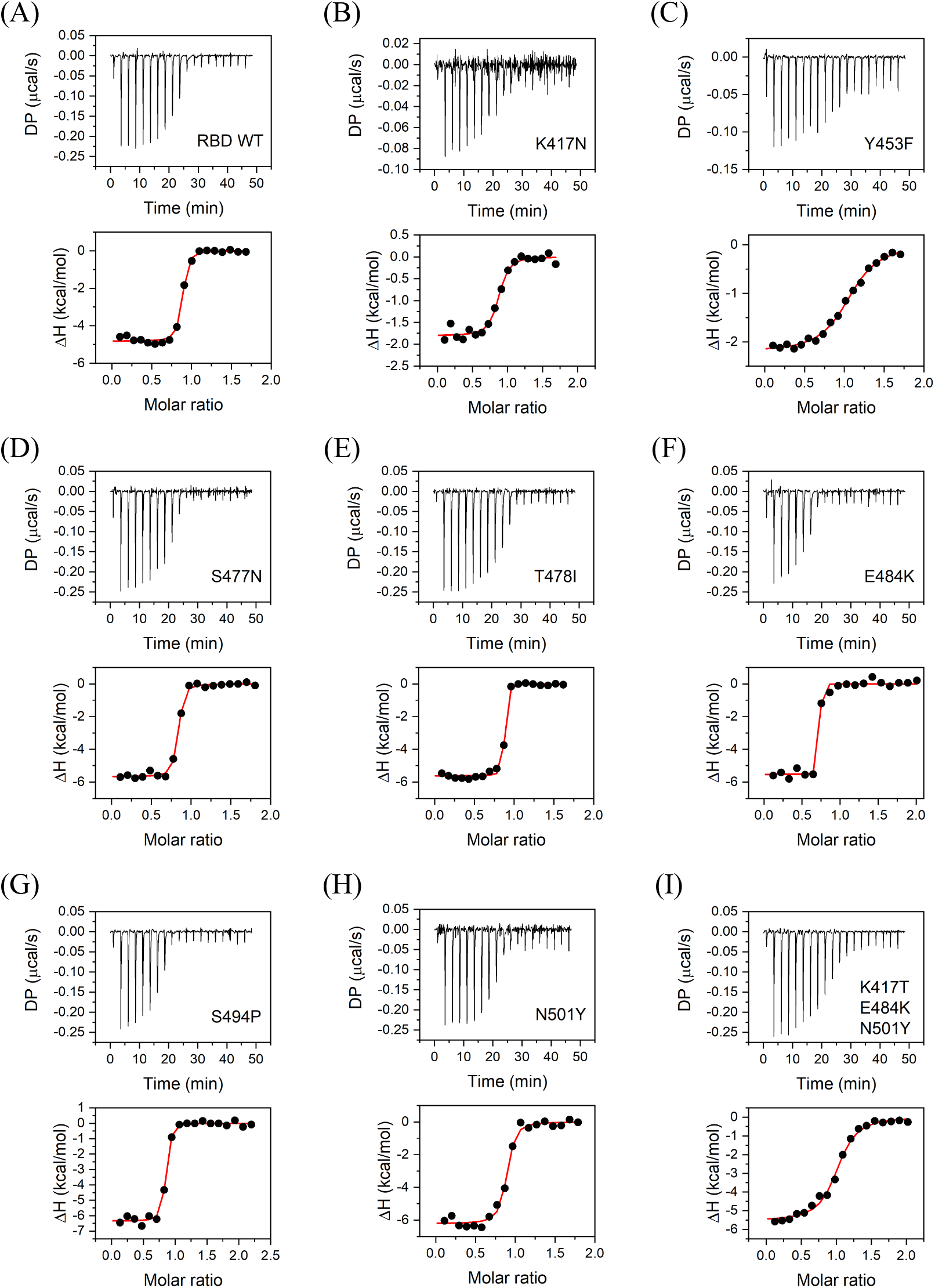
Binding of RBD and its variants to CC12.1 ScFv studied using ITC. Panels A-I show the data for the wild-type RBD, single amino acid mutations K417N, Y453F, S477N, T478I, E484K, S494P, N501Y, and for the triple mutant K417T/E483K/N501Y, respectively. Top panels show the raw thermograms, and the bottom panels show the fit to the integrated heat curve. Table 4 lists the interaction parameters from the data fit.

**Table 4.** Interaction parameters obtained from binding of RBD variants to CC12.1 ScFv probed by ITC. Errors on ΔG and -TΔS were calculated using error propagation formulae (72).

| **RBD variant** | **K<sub>d</sub>**<br>**(nM)** | **N** | **$\Delta H$**<br>**(kcal/mol)** | **$\Delta G$**<br>**(kcal/mol)** | **$-T\Delta S$**<br>**(kcal/mol)** |
| --- | --- | --- | --- | --- | --- |
| Wild-type | 28.0 ± 8.8 | 0.9 ± 0.0 | -4.8 ± 0.1 | -10.1 ± 0.2 | -5.3 ± 0.2 |
| K417N | 119 ± 50 | 0.8 ± 0.0 | -1.8 ± 0.1 | -9.3 ± 0.2 | -7.5 ± 0.3 |
| Y453F | 827 ± 146 | 1.0 ± 0.0 | -2.2 ± 0.1 | -8.2 ± 0.1 | -6.0 ± 0.1 |
| S477N | 25.6 ± 6.8 | 0.8 ± 0.0 | -5.7 ± 0.1 | -10.2 ± 0.2 | -4.5 ± 0.2 |
| T478I | 5.8 ± 3.4 | 0.8 ± 0.0 | -5.6 ± 0.1 | -11.0 ± 0.3 | -5.4 ± 0.4 |
| E484K | 1.7 ± 4.6 | 0.7 ± 0.0 | -5.5 ± 0.1 | -11.8 ± 1.6 | -6.2 ± 1.2 |
| S494P | 23.0 ± 7.7 | 0.8 ± 0.0 | -6.4 ± 0.1 | -10.2 ± 0.2 | -3.9 ± 0.2 |
| N501Y | 63 ± 22 | 0.9 ± 0.0 | -6.2 ± 0.2 | -9.7 ± 0.2 | -3.4 ± 0.3 |
| K417T/E484K/N501Y | 433 ± 95 | 1.0 ± 0.0 | -5.6 ± 0.2 | -8.5 ± 0.3 | -3.0 ± 0.2 |

## DISCUSSION

The reports of mutations in the spike protein of SARS-CoV-2 started to appear very soon after its emergence in Wuhan, China in December 2019. Most of the mutations in the RBD were noticed during the Fall of 2020, and variant B.1.1.7 with mutation N501Y (Alpha variant) became the first RBD variant to be labelled as a variant of concern (VOC) (https://www.who.int/en/activities/tracking-SARS-CoV-2-variants/). Since the beginning of 2021, there has been an emergence of numerous variants in different parts of the world. The rate of mutations in SARS-CoV-2 genome is lesser than that known for other RNA viruses like influenza and HIV (14-16). The prime reason for that is the ability of the virus to proof-read errors included by RNA-dependent RNA polymerase due to the presence of exonuclease activity (49). Even then, substantial number of variants occurring with high frequency were reported for this virus, with the list continuing to grow. This can be explained by the fact that the appearance of mutations is also dependent on the viral population size. Higher viral population will support greater number of mutants. As this virus has infected human population globally and has emerged as a pandemic, it is not surprising to encounter the different variants. The biggest concern with the emergence of the variants is the ineffectiveness of the currently developed vaccines, which are being used to vaccinate people, and the antibody-based therapeutics, which are approved for emergency use. There are reports of breakthrough infection in already vaccinated individuals (50) and also reports of decreased neutralization ability of vaccinated individuals’ serum against variants (36-38), which substantiate the need to study these variants in more detail.

The occurrence and persistence of a particular mutation in virus pool is dependent on a number of factors. In general, it is considered that mutations that provide a selective advantage to virus fitness are naturally selected. The most important characteristics of proteins that can affect virus fitness are its stability and activity. Mutations in viral proteins can have a neutral, stabilizing or destabilizing effect on protein stability. Stabilizing mutations offer a fitness advantage to the virus by increasing the proportion of correctly folded protein and increased resistance to protein degradation and aggregation inside the cell (51,52). Higher stability, similar to higher expression, can thus lead to higher infectivity through increased virus yield (53). Destabilizing mutations on the other hand do not offer fitness advantage and are mostly deleterious (54). They lead to lower virus yield with decreased infectivity. For example, in the case of hemagglutinin protein of Influenza virus, it has been shown that the higher stability variants provide fitness advantage and tend to persist longer (53). The mutation rate and the population size also affect the stability of the new variants. It has been shown that higher mutation rates and lower population size lead to the emergence of low stability variants and on the contrary low mutation rates and high population sizes lead to emergence of variants with higher stability (55). Our results show that the emerging variants are quite resistant to major stability changes (Table 2), despite the nature of the mutation (including many non-conservative mutations with differing physical properties) and multiple mutations accumulating in VOCs such as the Gamma variant.

Protein activity is another parameter that affects virus fitness. Unlike protein stability that can be impacted by mutations at many sites on protein, protein activity is controlled by a few key amino acid residues. In case of SARS-CoV-2 RBD, binding to ACE2 with a higher affinity provides a selective advantage towards virus fitness. Higher the affinity, higher the virus infectivity. The key residues of RBD which interact with ACE2 residues are shown in Fig. 7A. Most of the frequently occurring mutations in RBD (Fig. 1A) are at the binding interface with ACE2. Our data shows that the four mutations Y453F, S477N, T478I and S494P do not impact ACE2 binding. T478 and S494 are not part of the binding interface with ACE2 (Fig. 7A), and hence mutations T478I and S494P do not bring any change to the binding affinity. Y453 also does not form any polar contacts with ACE2. Its mutation to phenylalanine is a conservative mutation and thus does not show any difference in binding affinity. Two mutations K417N and E484K show a decrease in ACE2 binding affinity. K417 residue in SARS-CoV-2 RBD has previously been shown to be responsible for increase in binding efficiency of RBD towards ACE2 (46). K417 residue in SARS-CoV-2 RBD helps in bringing a more positive charge on the RBD surface which better interacts with the negatively charged residues on ACE2. Mutation of lysine to an uncharged amino acid asparagine decreases the positive charge on the surface thereby decreasing binding affinity towards ACE2. Similarly, E484, a negatively charged amino acid interacts with a positively charged amino acid K31 of ACE2. Mutation to lysine, a positively charged amino acid abolishes this interaction, explaining the decrease in the binding affinity. The other single site mutation N501Y corresponding to the Alpha variant and the triple mutant K417T/E484K/N501Y corresponding to the Gamma variant show increased binding affinity to ACE2. From this data, it appears that receptor binding is a big factor in driving SARS-CoV-2 variant evolution, but it is not the only driving force. Some of the RBM mutants are naturally selected even though they do not have increased affinity for ACE2.

**Figure 7.**
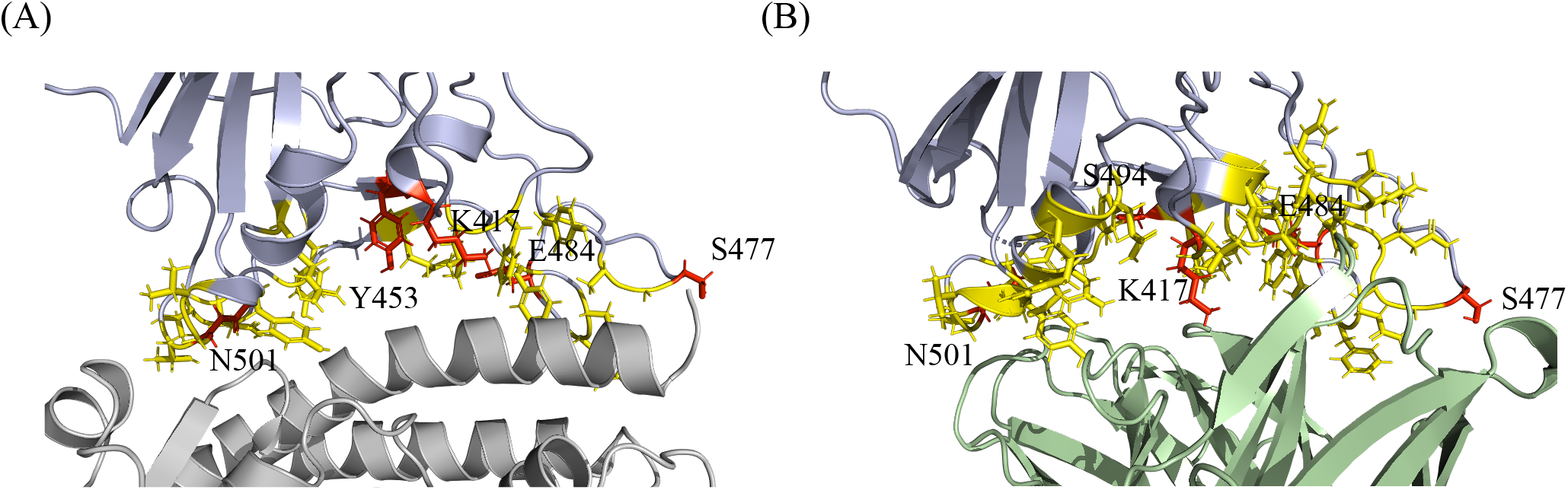
A model of SARS-CoV-2 RBD colored blue interacting with (A) ACE2 colored gray and (B) CC12.1 Fab colored green, showing the interface residues of RBD in yellow along with the side chains. The frequently mutating residues that are part of binding interface are colored red and are also shown with their side chains.

Neutral mutations which do not offer any advantage towards virus fitness and even deleterious mutations can get selected and become prevalent in the presence of other selection pressures like escape from human immune responses (56). Neutralizing antibodies are a key component of immune response against natural infection by viruses. Neutralizing antibodies can also be administered as recombinant monoclonal antibody therapeutics and as convalescent plasma to provide passive immunity. The circulating neutralizing antibodies thus provide a selection pressure that can drive the evolution of virus variants (57-59). Variants which can escape the neutralizing antibodies have a selective advantage and can persist and spread to other individuals. The effectiveness with which a variant can escape the neutralizing antibodies and become prevalent depends on the exposure of that variant to neutralizing antibodies. In the course of normal infection, the levels of antibody during the virus transmission phase are negligible. Thus, the transmitting viruses do not have exposure to neutralizing antibodies and are not considered to contribute to spread of variants with antibody escape potential (60). But reinfection in people with weak immunity increases the chances of exposure of virus to neutralizing antibodies elicited during first infection and contribute to selection of variants with escape potential (61-63). Similarly, convalescent plasma therapy, monoclonal antibody therapy (especially single antibody) and less immunogenic vaccine candidates can increase the chances of variants coming in contact with neutralizing antibodies and drive the emergence of variants that can escape these antibodies (64).

In this study, we tested the potential of the most prevalent variants to escape CC12.1 antibody, a naturally elicited human antibody as a response to SARS-CoV-2 infection (47). It belongs to a class of antibodies that target RBD and is the most abundant class of antibodies naturally produced by humans (20,48,65). Antibodies belonging to this class are encoded by the VH3-53 gene segment and represents a set of neutralizing antibodies that bind to the RBD epitope that overlaps with ACE2 binding epitope (Fig. 1) (20). These antibodies are characterized by short H3 CDR and can bind the RBD in “up” conformation. Nevertheless, among the RBD targeting antibodies, these are the most abundant and commonly found antibodies and represent the general antibody response to SARS-CoV-2 infection in humans (48). Our study examined which of the most prevalent variants have the ability to escape natural immune responses, but it should also be considered that the neutralizing antibody response towards an infection is varied with several neutralizing antibodies generated against different epitopes of spike protein (66,67). Also, the natural antibody responses can vary from individual to individual. Even then, neutralizing antibodies against RBD represents the most potent and widely used countermeasures against COVID-19 (68). The RBD-CC12.1 binding interface is shown in Fig. 7B. Most of the frequently mutating residues we examined are also part of the binding interface, which implies that the mutations are quite likely to impact CC12.1 binding. Our data shows that only 4 out of the 8 mutants (K417N, Y453F, N501Y and the triple mutant K417T/E484K/N501Y) we examined bind to CC12.1 with weaker affinity (or higher K_d_) (Table 4). The single site mutant Y453F showed the weakest binding or the highest potential to escape CC12.1 ScFv. Interestingly, Y453 is not a part of the binding interface with CC12.1, implying that the residues which do not directly interact with its binding partner can also impact binding after mutation. Other 2 mutants S477N and S494P did not show any difference in their binding affinity to CC12.1. Interestingly, E484K and T478I bind with a higher affinity (or lower K_d_) to CC12.1 (Table 4). These results show that the neutralizing antibody response is one of the driving forces for natural selection of RBD variants and should be considered in closer detail. It should also be considered that the results presented here are only representing the escape towards one class of antibody, and other antibodies that are naturally occurring or administered passively would also have an impact on the emergence of these variants. For example, although E484K and T478I mutations may not escape CC12.1 (Table 4), they may escape other classes of antibodies (69,70). These results also stress on the need to use viral sequencing to find the variant that has infected a patient and use of those antibodies to treat patients against which the variant does not show escape potential. To achieve this, different therapeutic neutralizing antibodies that target different epitopes on RBD should be developed, and a cocktail antibody drug may work better against new variants.

In **Summary**, we have examined the various factors that might be contributing to the natural selection of most frequent SARS-CoV-2 RBD variants, in particular protein expression, stability, activity in terms binding to ACE2 and antibody escape potential in terms of binding to a human neutralizing antibody. Table 5 summarizes our observations with red color indicating changes that favor natural selection of variants. These results show that multiple factors contribute to the natural selection of variants and should be considered when evaluating any future variants. For example, the triple mutant K417T/E484K/N501Y (Gamma variant) poses a serious threat with many factors, increased protein expression, increased activity and increased antibody escape potential favoring its emergence and persistence. Followed by the Gamma variant, Alpha variant N501Y has the most favorable biophysical parameters in terms of increased affinity towards ACE2 and increased escape potential (Table 5). In the case of variants harboring multiple mutations, each mutation might be playing a specific role in virus survival. It can be either stronger binding to ACE2, increased expression, or escape from neutralizing antibodies. For example, when the data in Table 5 is compared between the N501Y (Alpha variant), the triple mutant K417T/E484K/N501Y (Gamma variant) and the Wild-type, increase in ACE2 binding might be significantly determined by the N501Y mutation, since no significant differences were observed in the K_d_ of ACE2 binding between the N501Y and K417T/E484K/N501Y variants. The effect of the other two mutations K417T and E484K in the Gamma variant might be to increase its expression and/or to increase immune escape potential. E484K by itself do not have increased expression or do not escape from CC12.1 compared to the wild-type (Table 5). In fact, E484K binds to CC12.1 with higher affinity, but it is quite likely that this mutation has evolved to show increased immune escape against other neutralizing antibodies (69,70). Out of the 8 variants for which complete biophysical data is available in Table 5, only two (N501Y (Alpha variant) and the triple mutant K417T/E484K/N501Y (Gamma variant)) correspond to the variants of concern (VOCs) and the other 6 are classified as variants of interest (VOI). The data suggests that VOCs are evolving by maximizing the biophysical fitness parameters that determine the virus survival. We hope that such fundamental understanding of the physical parameters determining the natural selection of variants will help in designing better countermeasures like new vaccine candidates and antibody therapeutics that work against emerging variants.

**Table 5.** Comparison of protein expression, stability, binding to ACE2, and binding to CC12.1 of RBD variants. Red colored values indicate parameters favoring the natural selection of variants.

| RBD variant | Expression | T<sub>m</sub> (°C) | K<sub>d</sub> (nM)<br>(ACE2 binding) | K<sub>d</sub> (nM)<br>(CC12.1 binding) |
| --- | --- | --- | --- | --- |
| Wild-type | - | 56.1 ± 0.7 | 10.0 ± 3.1 | 28.0 ± 8.8 |
| K417N | No change | 56.9 ± 0.3 | 94.1 ± 9.5 | 119 ± 50 |
| Y453F | Increase | 54.8 ± 0.4 | 11.4 ± 1.7 | 827 ± 146 |
| S477N | Increase | 55.8 ± 0.3 | 5.0 ± 1.4 | 25.6 ± 6.8 |
| T478I | Decrease | 52.3 ± 0.2 | 16.7 ± 5.6 | 5.8 ± 3.4 |
| E484K | Decrease | 52.3 ± 0.5 | 51.5 ± 8.2 | 1.7 ± 4.6 |
| S494P | Increase | 55.3 ± 0.4 | 13.8 ± 3.5 | 23.0 ± 7.7 |
| N501Y | No Change | 55.1 ± 0.3 | 3.0 ± 2.1 | 63 ± 22 |
| K417T/E484K/N501Y | Increase | 56.5 ± 0.2 | 1.6 ± 1.5 | 433 ± 95 |

## EXPERIMENTAL PROCEDURES

### Cloning and expression of RBD, RBD mutants, ACE2 and CC12.1 ScFv

Amino acid sequences of SARS-CoV-2 RBD and human-ACE2 protein were obtained from Uniprot (Uniprot ID-P0DTC2 and Q9BYF1 respectively). The sequence of heavy and light chain variable regions (V_H_ and V_L_ respectively) of CC12.1 neutralizing antibody was obtained from RCSB-PDB (PDB ID 6XC2). The single chain variable fragment (ScFv) for CC12.1 was designed as V_H_-(GGGGS)_3_-V_L_. The protein sequences were codon-optimized for expression in human cells and synthesized by Twist Biosciences. The RBD variants were generated by site directed mutagenesis using mutagenic primers. The synthesized and mutated genes were cloned into pcDNA 3.4 Topo vector, modified by including a signal sequence of human immunoglobulin heavy chain, His-Tag and SUMOstar protein. The expression vectors were transfected transiently into Expi293 (modified HEK293) cells using polyethylenimine (PEI) and the protein was recovered from the culture supernatant after 5 days. The expression levels for the proteins were compared after running the culture supernatants on SDS-PAGE, staining with Coomassie blue R-250 dye, and quantifying the band intensities corresponding to the target protein using the ImageLab software from Bio-Rad.

### Protein Purification

The supernatant of the culture was filtered through 0.22 μm filter and purified using Nickel-nitrilotriacetic acid (Ni-NTA) chromatography. The eluted protein was dialyzed in buffer containing 50 mM of Tris-HCl, 20 mM NaCl, pH 8.0 and digested using SUMOstar protease overnight to cleave the target protein from SUMOstar and His-Tag. The digested protein was passed again through the Ni-NTA column to obtain the untagged target protein in the flow-through. All proteins were dialyzed in buffer containing 50 mM Sodium Phosphate, 20 mM NaCl, pH 7.0.

### Isothermal titration calorimetry (ITC)

ITC experiments were performed using MicroCal-PEAQ-ITC instrument from Malvern in buffer containing 50 mM Sodium Phosphate, 20 mM NaCl, pH 7.0 at 20° C. For ACE2-RBD interactions, ACE2 at a concentration of 12 μM was taken in the cell and RBD variants at concentration of 120 μM were taken in the syringe. For RBD-CC12.1 ScFv interactions, RBD or its variants were taken in the cell at a concentration of 20 μM and CC12.1 ScFv was taken in the syringe at 200 μM. The syringe contents were titrated into the cell as 18 injections of 2 μl each with 150 s spacing between the injections. ITC data analysis was done using MicroCal PEAQ-ITC Analysis Software from Malvern.

### Circular dichroism (CD) spectroscopy

CD spectra of RBD variants were recorded on an Applied Photophysics Chirascan Plus spectrometer at a protein concentration of 5 μM in buffer containing 10 mM Sodium Phosphate, 4 mM NaCl, pH 7.0 in a 1 mm pathlength cuvette. The data was collected at an interval of 1 nm and averaged for 2 s at each wavelength.

The thermal melts for RBD variants were recorded at a protein concentration of 20 μM in buffer containing 50 mM Sodium Phosphate, 20 mM NaCl, pH 7.0 in 0.5 mm pathlength cuvette. The temperature scan rate was 1° C/min. CD signal at 222 nm averaged for 2 s was plotted against the temperature and fitted to a two-state unfolding model using the equation (71),

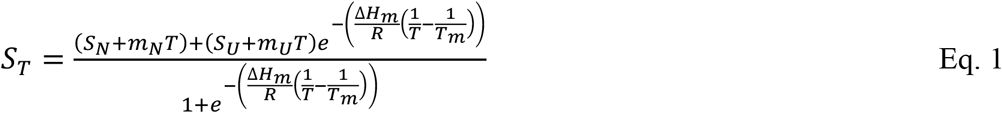

where S_T_ is the measured signal as a function of temperature T, S_N_ and S_U_ are the signals corresponding to the native and unfolded baselines, m_N_ and m_U_ are the slopes of linear dependence of S_N_ and S_U_, ΔH_m_ is the enthalpy change at T_m_, R is the universal gas constant and T is the absolute temperature in Kelvin.

## Abbreviations

ACE2: angiotensin converting enzyme 2;
CC12.1: neutralizing antibody from COVID-19 survivors;
CD: circular dichroism;
COVID-19: coronavirus disease 2019;
HEK: human embryonic kidney;
ITC: isothermal titration calorimetry;
MERS-CoV: Middle East respiratory syndrome coronavirus;
Ni-NTA: nickel nitrilotriacetic acid;
PEI: polyethylenimine;
RBD: receptor binding domain;
RBM: receptor binding motif;
SARS-CoV: severe acute respiratory syndrome coronavirus;
SARS-CoV-2: severe acute respiratory syndrome coronavirus 2;
ScFv: single chain variable fragment;
Tm: midpoint temperature of thermal denaturation;
VOC: variant of concern;
VOI: variant of interest.

## FIGURE LEGENDS

**Figure 1**. Structures of SARS-CoV-2 RBD (colored blue) interacting with (A) ACE2 (colored gray; PDB ID 6m0j) and (B) CC12.1 Fab (colored green; PDB ID 6xc2), showing the most frequently mutating residues in RBD-K417, N439, Y453, S477, T478, E484, S494 and N501 (colored red). RBM is shown in yellow color. The single mutants of RBD used in this study were K417N, N439K, Y453F, S477N, T478I, E484K, S494P and N501Y (Alpha variant). A double mutant (E484K/N501Y) and triple mutants corresponding to Beta variant (K417N/E484K/N501Y) and Gamma variant (K417T/E484K/N501Y) were also used. The position of Y453 is not visible in the surface view of RBD interacting with CC12.1 Fab as it is buried at the interface.

**Figure 2**. Comparison of relative expression of RBD and its mutants. (A) SDS-PAGE showing relative amounts of expressed RBD and its mutants. M represents molecular weight markers (From top to bottom: 180, 130, 100, 70, 55, 35 and 25 kDa) (B) Relative expression of mutants quantified from the band intensities in SDS-PAGE in panel A.

**Figure 3**. (A) Purified RBD and its mutants (B) ACE2 and CC12.1 ScFv. M represents molecular weight marker (From top to bottom: 180, 130, 100, 70, 55, 35, 25, 15 and 10 kDa). (C) Comparison of secondary structures of RBD and its mutants using far-UV CD spectroscopy. Table 1 lists the proportion of various secondary structures when the spectra were deconvoluted using BeStSel software.

**Figure 4**. Thermal denaturation melts of RBD and its mutants obtained using far-UV CD spectroscopy. Panels A-I show the data for the wild-type RBD, single amino acid mutations K417N, Y453F, S477N, T478I, E484K, S494P, N501Y, and for the triple mutant K417T/E483K/N501Y, respectively. The solid lines show the fits to a 2-state unfolding equation (Eq. 1 in Methods section). Table 2 lists the T_m_ (midpoint meting temperature) and the DH (enthalpy change at Tm) values of RBD variants.

**Figure 5**. Binding of RBD and its variants to ACE2 studied using ITC. Panels A-I show the data for the wild-type RBD, single amino acid mutations K417N, Y453F, S477N, T478I, E484K, S494P, N501Y, and for the triple mutant K417T/E483K/N501Y, respectively. Top panels show the raw thermograms, and the bottom panels show the fit to the integrated heat curve. Table 3 lists the interaction parameters from the data fit.

**Figure 6**. Binding of RBD and its variants to CC12.1 ScFv studied using ITC. Panels A-I show the data for the wild-type RBD, single amino acid mutations K417N, Y453F, S477N, T478I, E484K, S494P, N501Y, and for the triple mutant K417T/E483K/N501Y, respectively. Top panels show the raw thermograms, and the bottom panels show the fit to the integrated heat curve. Table 4 lists the interaction parameters from the data fit.

**Figure 7**. A model of SARS-CoV-2 RBD colored blue interacting with (A) ACE2 colored gray and (B) CC12.1 Fab colored green, showing the interface residues of RBD in yellow along with the side chains. The frequently mutating residues that are part of binding interface are colored red and are also shown with their side chains.

